# Quantitative resistance loci to southern rust mapped in a temperate maize diversity panel

**DOI:** 10.1101/2021.04.02.438220

**Authors:** Guangchao Sun, Ravi V. Mural, Jonathan D. Turkus, James C. Schnable

## Abstract

Southern rust is a severe foliar disease of maize resulting from infection with the obligate biotrophic fungus, *Puccinia polysora*. The disease reduces photosynthetic productivity which reduces yields with the greatest yield losses (up to 50%) associated with earlier onset infections. *Puccinia polysora* urediniospores overwinter only in tropical and subtropical regions but cause outbreaks when environmental conditions favor initial infection. Increased temperatures and humidity during the growing season, combined with an increased frequency of moderate winters are likely to increase the frequency of severe southern rust outbreaks in the US corn belt. In summer 2020, a severe outbreak of Southern Rust was observed in eastern Nebraska (NE), USA. Disease incidence severity showed significant variation among maize genotypes. A replicated maize association panel planted in Lincoln, NE was scored for disease severity. Genome wide association studies identified four loci associated with significant quantitative variation in disease severity which were associated with candidate genes with plausible links to quantitative disease resistance and a transcriptome wide association study conducted identified additional associated genes. Together these results indicate substantial diversity in resistance to southern rust exists among current temperate adapted maize germplasm, including several candidate loci which may explain observed variation in resistance to southern rust.

## Introduction

Southern rust is a foliar disease resulting from infection with the fungus *Puccinia polysora* which infects corn (*Zea mays*) throughout the world although the greatest yield losses are observed in tropical and semitropical regions. In the north central United States, southern rust outbreaks are infrequent as the spore cannot overwinter in the region. However, when an outbreak occurs, sometimes following a moderate winter, yield losses can be substantial as temperate maize germplasm predominantly lacks both qualitative and quantitative resistance to southern rust^1^. Yield losses of 138 million bushels were attributed to an outbreak of southern rust in the northern United States and Canada in 2016, and 93 million bushels of lost yield in the same region in 2017^2^. Yet outbreaks of southern rust can be much worse. In the 1950s southern rust reached Africa where the maize varieties currently being grown lacked resistance to the pathogen and yield losses of 50% were observed across large regions^3^. In controlled inoculation studies, maize yields were reduced 39% in response to inoculation with *Puccinia polysora* in Maryland^4^. Comparisons of near isogenic resistant and susceptible hybrids in Mississippi indicated southern rust can reduce yields by up to 45%^5^ with the severity of the losses depending on how early in its life cycle the corn field is exposed to southern rust infection.

The first reported observation of southern rust was on *Tripsacum dactyloides*, a wild relative of maize, outside Auburn, Alabama in 1891^6^. However, the pathogen was likely widespread in central and southern America by 1879^7^. *Puccinia polysora* is cold sensitive and cannot overwinter in the USA. The severity of southern rust outbreaks in the northern USA depends on how far south the urediniospores of the pathogen died off in the previous winter, how quickly urediniospores produced in the south blow northwards, and how rapidly the urediniospores are able to initiate infection based on the environmental conditions^8^. Given changes in the severity of winters in the temperate United States, it is likely that southern rust outbreaks in the northern USA will become more common and more severe in coming years.

Southern rust can be controlled with a range of fungicides^8^. However, the most economical and sustainable approach to mitigating yield losses is the development and commercialization of resistant maize lines. A number of resistance loci have been reported in both temperate and tropical germplasm, with many mapped resistance loci clustering on the short arm of chromosome 10, some or all of which may represent alleles of the same locus^1,9^. The *RPP9* locus on chromosome 10 was originally identified in Boesman Yellow Flint, a maize line collected in South Africa^10^. In the 1990s commercial hybrids were released in the USA with resistance to a prevalent strain of southern rust^11^. By 2009, southern rust infections were reported in lines carrying the *RPP9* locus after successfully controlling southern rust for 30 years, suggesting the breakdown of resistance conferred by this locus^1,12,13^. The potential for the breakdown of resistance is a major downside of qualitative resistant genes. In contrast, outcome of the majority of host-pathogen interactions are determined by quantitative resistance mechanisms which are less susceptible to breakdown, but also less amenable to genetic investigation and incorporation into breeding programs^14^.

A number of reports of quantitative resistance to southern rust exist in the literature. Holland and co-workers mapped quantitative resistance in F2 derived F3 populations. In addition to the widely reported large effect loci on the short arm of chromosome 10, two smaller effect QTLs (quantitative trait loci) on chromosomes 3 and 4 with each explaining approximately 14% of total variation in southern rust resistance were also identified^11^. In addition, Jiang and co-workers reported QTLs on chromosomes 3, 4 and 9^15^; Brunelli and co-workers identified a QTL on chromosomes 9^16^; Jines and co-workers mapped QTLs on chromosomes 4, 8, 9 and 10^17^; Brewbaker and co-workers mapped a QTL on chromosome 6^1^ and in 2013, Wanlayaporn *et al*., reported six QTLs for resistance on chromosomes 1, 2, 5, 6, 9 and 10^18^. These studies in aggregate demonstrate that both qualitative and quantitative southern rust resistance loci exist within maize germplasm. However, they primarily utilized biparental populations which, while increasing the power to detect segregating variation, provide a limited amount of insight into the overall genetic architecture of quantitative resistance to southern rust in temperate maize.

A severe outbreak of southern rust in eastern Nebraska during the 2020 growing season^19,20^ provided an opportunity to screen a preexisting maize experiment constituting 752 diverse genotypes grown in a replicated complete block design for quantitative severity of southern rust infection. These genotypes were primarily drawn from temperate germplasm^21^ and hence previously reported large effect qualitative resistance loci such as *RPP9* which are near exclusively drawn from tropical germplasm pools are unlikely to be segregating in this population^1^. Employing a genotype dataset consisted of 2.7 million SNP markers with a minor allele frequency >0.1 generated from a combination of the published maize hapmap3^22^, deep resequencing data for 509 Wisconsin Diversity Panel^23^ and RNAseq data^21,24^ multiple loci associated with quantitative resistance to southern rust were identified through a combination of genome wide association and transcriptome wide association. The latter utilized published expression data collected from seedlings^21,24^. In aggregate, these results indicate that, while quantitative resistance to southern rust is a polygenic trait with a complex genetic architecture, loci contributing to variation are present and detectable in existing temperate maize germplasm.

## Results

### Observed distribution of southern rust severity in a maize diversity panel

A set of 752 lines drawn from the predominantly temperate Wisconsin Diversity Panel^21,26^ were grown in a replicated design in a farm near Lincoln, NE (see methods). On August 14^th^, southern corn rust pustules were observed on the leaf surfaces of corn plants of multiple genotypes at the R1-R4 stage (Figure 1A). Based on the life cycle of southern corn rust and southern corn rust tracker^20^, the infection was likely started in Mid July during which the environment at the Nebraska field site was humid with frequent rainfalls, and warm winds from south may have carried a considerate number of spores into the region. On August 19^th^ of 2020, disease severity scores were obtained for the field by three individual scorers, using a published disease severity reference ranging from 0 (no disease) to 4 (high susceptibility)^27^. Among the 1,680 plots of the original field layout, 286 were severely lodged or had already senesced and 1,398 plots, representing 689 genotypes were successfully scored. Each scorer was assigned partially overlapping sets of plots resulting in 2,065 unique disease severity score data points being collected (Figure S1). Individual scorers produced distinct medians and standard deviations for disease severity scores (Figure 1B). To confirm these ranges did not reflect spatial variation in disease severity among plots assigned to only one scorer, a comparison was conducted among plots assigned to all three scorers, and similar differences in median and standard deviation were observed (Figure 1C). This significant human bias in the assignment of disease severity scores was consistent, yet at the same time pairwise comparisons among scorer showed consistent correlations in the disease severity scores assigned to individual plots (Figure 1D).

**Figure 1.**
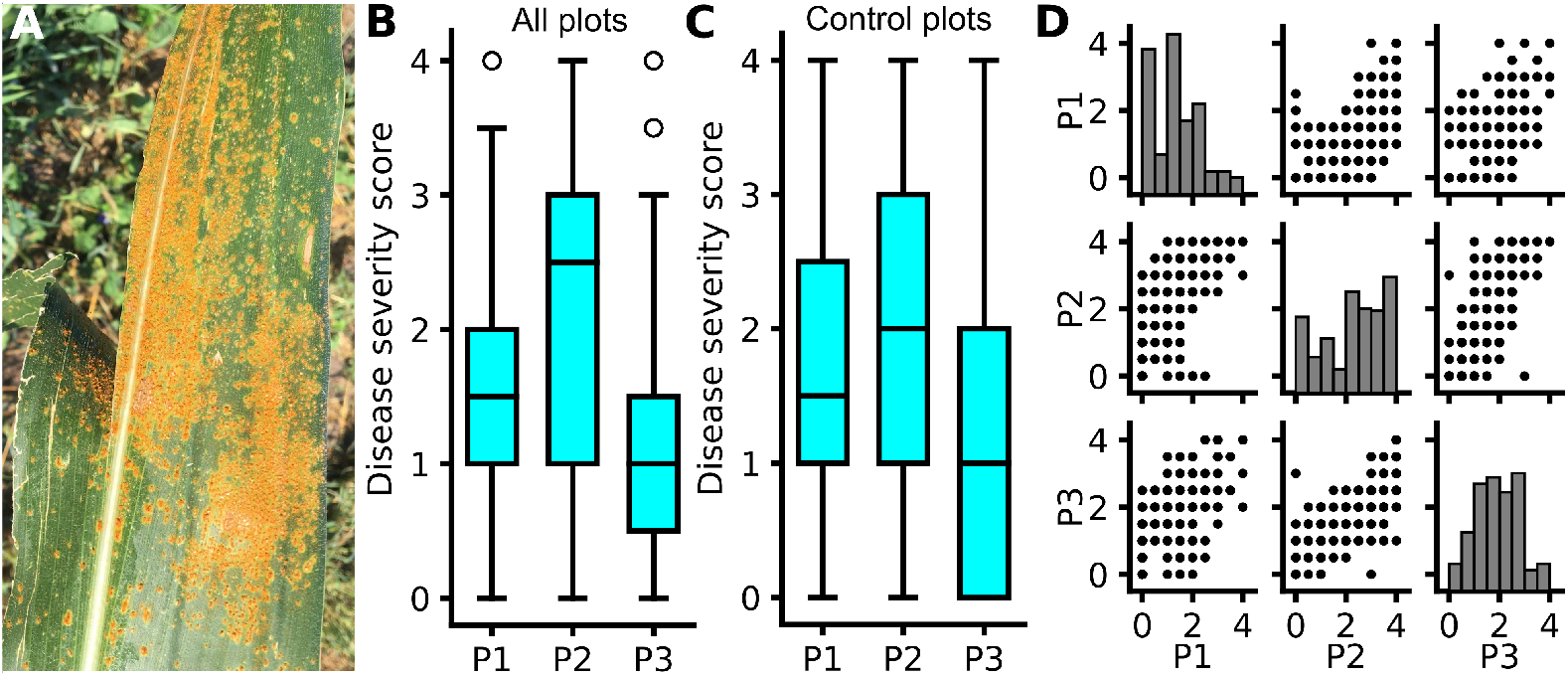
Observed southern rust disease severity across an association panel in 2020. A) An example of an infected leaf from a susceptible maize line of which disease severity scored as 4. B) Distribution of disease severity scores recorded by three different observers, P1-P3, across the experiment (1505 total plots). Not all plots were scored by each individual. C) Distribution of of disease severity scores recorded by the same three observers on a set of 272 plots which were independently scored by each of the three individuals. C) Pair-wise comparison of the disease severity scored for the same plots by three scorers. Score distribution were shown in the diagonal.

Correlations between the disease severity scores assigned to the same plots by P1 and P2 as well as P2 and P3 were greater than the correlations between disease severity scores assigned by P1 and P3. The P2 scores were more widely distributed across the entire scoring range. Using data on plots scored by all three individuals, models were trained to predict P2 scores using either the P1 or P3 scores, combined with spatial data, to predict the disease severity score which would have been assigned by P2. The R^2^ of the regression line between the raw disease severity scored by P1 and P2 was 0.443 and that between P3 and P2 was 0.555 respectively. Corrected scores for P1 and P3 exhibited significant shrinkage towards the regression line but substantially corrected for differences in median and standard deviation in the raw scores (Figure 2A & B). Raw disease severity scores exhibited a heritability (H^2^) of 0.45. However, after correction for scorer specific biases in median and standard deviation of results, corrected disease severity scores exhibited a heritability of H^2^ = 0.56. The 689 lines scored as part of this field included seven lines rated either 0 or 1 in a southern rust resistance screen of 1,890 diverse maize associations in Illinois using controlled inoculations of *Puccinia polysora*^28^. Among overlapping lines identified as resistant (0 or 1) in controlled inoculation studies, the average best unbiased linear predictor (BLUP) assigned to a line in this study was 1.2, with a maximum of 1.7 (Table S1).

**Figure 2.**
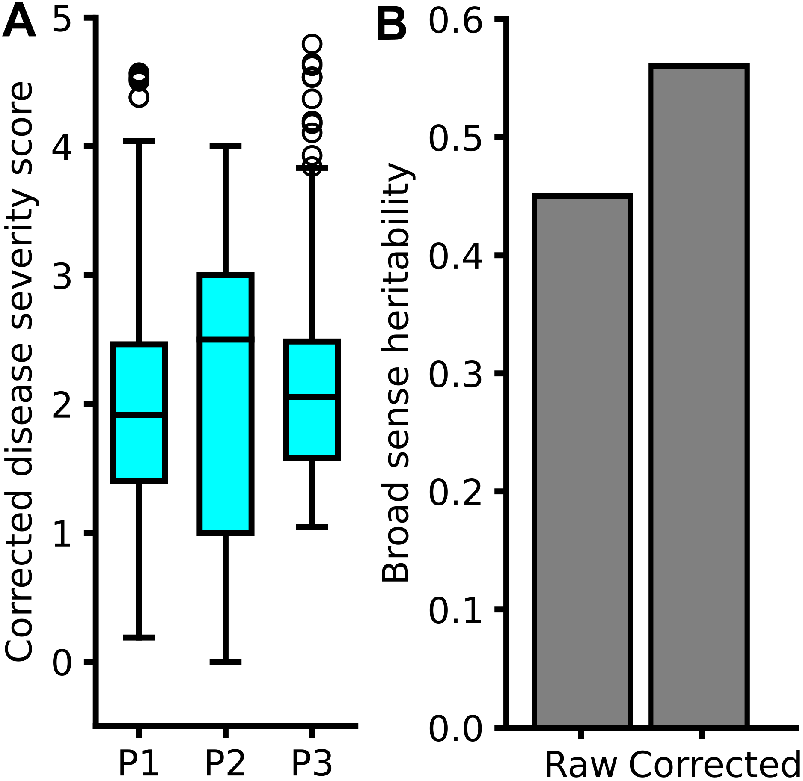
Correcting for individual patterns in disease severity scores. A) Distribution of disease severity scores after the raw scores for scorer1 (P1) and scorer3 (P3) were replaced by predicted scored generated using a model trained to predict the values scorer2 with both human error and spatial effects incorporated (P2 would assign to a given plot using the scores assigned by P1 or P3). B) Mostly higher broad sense heritability was observed for scorer-bias-corrected disease severity scores (H^2^=0.56) than for the raw values (H^2^=0.45).

### Genomic loci linked to variation in southern corn rust disease severity

The heritability of southern rust severity scores amongst the lines scored as part of this experiment suggested it might be possible to identify individual genetic loci which contribute to the observed severity of southern rust infections. Using a set of SNP markers generated from a combination of public resequencing experiments, maize hapmap and RNA sequencing datasets^21–24^, Genome Wide Association Studies (GWAS) were conducted to detect potential genomic loci or features significantly associated with variation in southern rust severity. A total of six distinct peaks on chromosomes 2, 4 (two peaks), 5, 6, and 7 which passed a suggestive p-value cutoff of 10e^-5^ were identified using GEMMA^25^. The quantile-quantile plot for this analysis showed significant deviation of p values from the expected null model suggesting true positive signals detected by GWAS (Figure S3). Of the six suggestive peaks, three were validated in *geq* 10% of FarmCPU resampling runs (Chr2:231,271,050, Chr4:78,851,667, Chr4: 173,863,109), and a fourth signal was detected on (Chr6:169,030,253) which did not correspond to a suggestive peak in the GEMMA results (Figure 3A). The absolute values of estimated effect sizes for these four loci ranged from 0.122 to 0.167 3B) corresponding to an estimated difference in disease severity score between inbreds homozygous for different alleles of between 0.244 and 0.334. Distributions of observed disease severity scores were notably different for each of the four loci, with the alternative alleles on chromosomes 2 and 6 associated with a decrease in southern rust severity and the alternative alleles of the two loci on chromosome 4 associated with an increase in southern rust severity (Figure 3C).

**Figure 3.**
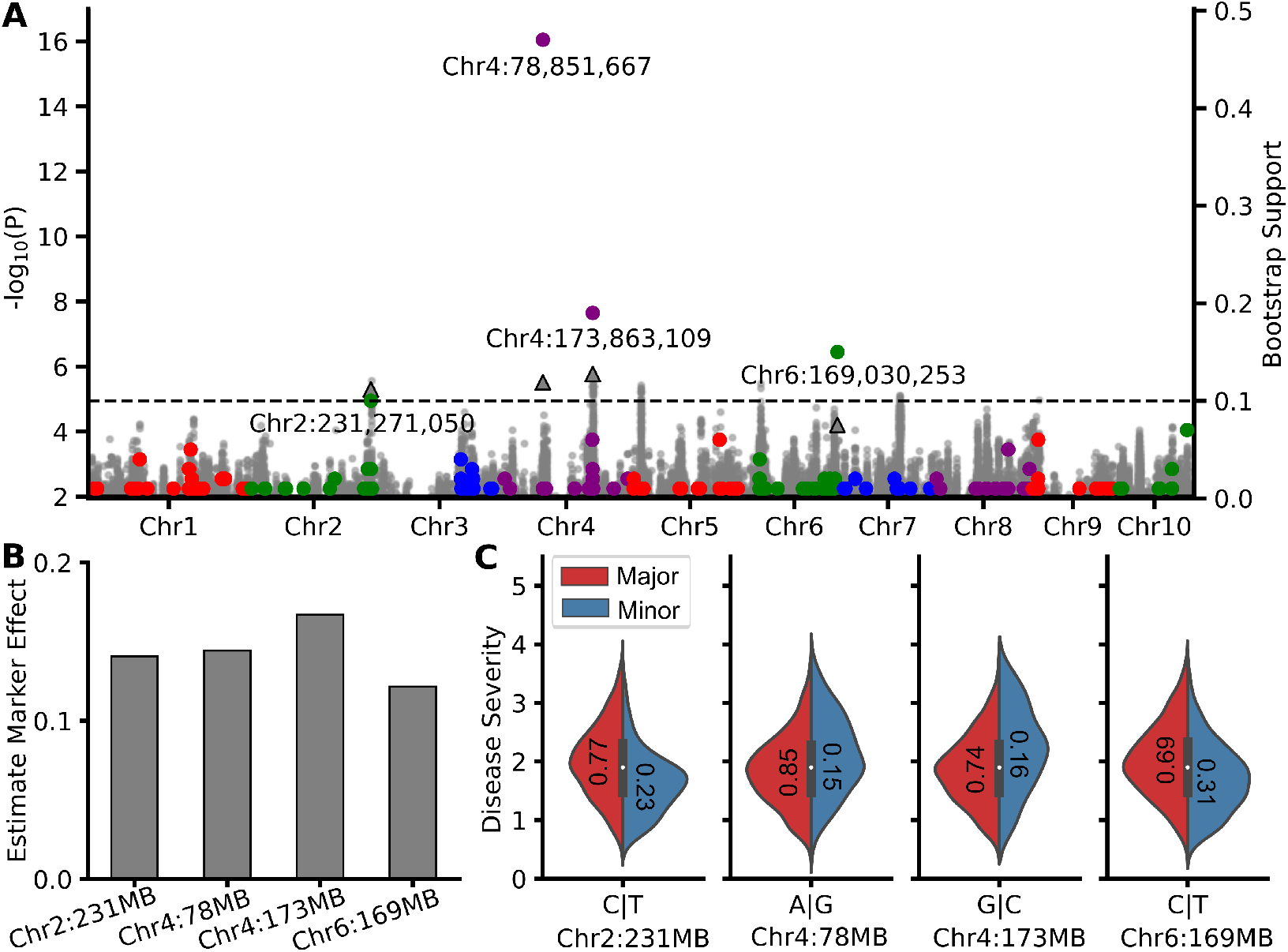
Results of a genome wide association study for southern rust resistance in temperate maize. A) Distribution across the genome for the statistical significance for the association with between individual SNPs and disease severity scores calculated using the MLM based GWAS algorithm implemented in GEMMA^25^ gray points and left y-axis. Genomic locations of significant marker/ disease severity associations identified in at least one out one hundred resampling FarmCPU GWAS analyses conducted, and the proportion of analyses in which a given marker was statistically significant (colored circles and right y-axis). For the ten markers which were statistically significant in at least 10% of FarmCPU resampling runs, the statistical significance assigned to the same marker by GEMMA is indicated by a black triangle. Dashed gray line indicates a cutoff of 10% for resampling support and a GEMMA p-value of 10e^-5^. B) Estimated absolute marker effect sizes for the four markers which were statistically significant in at least 10% of FarmCPU resampling runs. C) Distributions of observed distributions of disease severity scores among inbreds carrying the reference (red) and alternate (blue) alleles for each of the four markers which were statistically significant in at least 10% of FarmCPU resampling runs, numbers in the plots were allele frequencies for major and minor alleles respectively.

Patterns of linkage disequilibrium vary throughout the maize genome can create challenges for linking trait associated SNPs to candidate genes. Each of the four trait associated SNPs was within one megabase up- and downstream of a large number of annotated gene models. Except for the second FarmCPU significant SNP on chromosome 4 at 173,863,109 that is within a gene Zm00001d051893, the distance to the first nearby genes for the FarmCPU significant marker on chromosome 2 at 231,271,050 bp was 30.8 kb, for the FarmCPU significant marker on chromosome 4 at 78,851,667 bp was 32.2 kb and for the FarmCPU significant marker on chromosome 6 at 169,030,253 was 1.6 kb respectively (Table S2). Markers within one megabase up and downstream of each of the four trait associated SNPs identified above were screened for linkage with the initially identified causal SNP. This analysis identified linked SNPs (R^2^ ≥ 0.8) for the significant SNP 2:231,271,050 spanning a 27 kb window where no gene resides, for the significant SNP 4:78,851,667 spanning a 1.4 kb window where no gene was located, for the significant SNP 4:173,863,109 spanning a 4.1 kb window where gene Zm00001d051893 resides and for the significant SNP 6:169,030,284 spanning a 0.1 kb window where no gene resides (Table S2).

It is also the case in maize that causal polymorphisms can exist in upstream regulatory regions which are only partially linked or even unlinked to the coding sequences the genes regulate. Noteable examples include gene Teosinte Branched1 (*Tb1*) where the polymorphism responsible for the suppression of axillary branching is associated with a long range cis-regulatory element 60 kb away where a Hopscotch insertion leading to variable *Tb1* transcription level^29, 30^; Vegetative to generative transition1 (*Vgt1*), a major QTL regulating flowering time through target gene *ZmRap2.7* expression regulation, was mapped to 70 kb away from the target gene^31,32^. Using public expression data from maize seedlings, we conducted eQTL analysis to identify cases where the trait associated SNP or another SNP in high linkage disequilibrium with the trait associated SNP (LD ≥ 0.8) was, in turn, significantly associated with variation of a gene, either inside or outside of the mapping interval defined by linkage disequilibrium. A total of twelve potential regulatory signals were linked with the four trait associated SNPs either directly, or indirectly via a SNP in high LD with the trait associated SNP. The expression levels of a single nearby gene, Zm00001d007424 was significantly associated with the trait associated SNP on chromosome 2 (Figure 4A & Figure S4A). The first trait associated SNP on chromosome 4 (at 79 MB) was directly associated with the expression levels of three genes Zm00001d050283, Zm00001d050284 and Zm00001d050293 (Figure 4B & Figure S4B-D). The second trait associated SNP on chromosome 4 (at 174 MB) was directly associated with the expression levels of two genes Zm00001d051914 and Zm00001d051893 and indirectly associated with the expression levels of two others Zm00001d051869 and Zm00001d05188 (Figure 4C & Figure S4E-F). The trait associated SNP on chromosome 6 was directly associated with the expression levels of two genes Zm00001d039039 and Zm00001d039020 and indirectly associated with the expression of an additional two genes Zm00001d039004 and Zm00001d039043 (Figure 4D & Figure S4G-I). For those genes whose expression variation directly associated with the four trait associated SNPs, the we individuals carrying different alleles did present a consistent correlation between expression level and disease severity (Figure S4)

**Figure 4.**
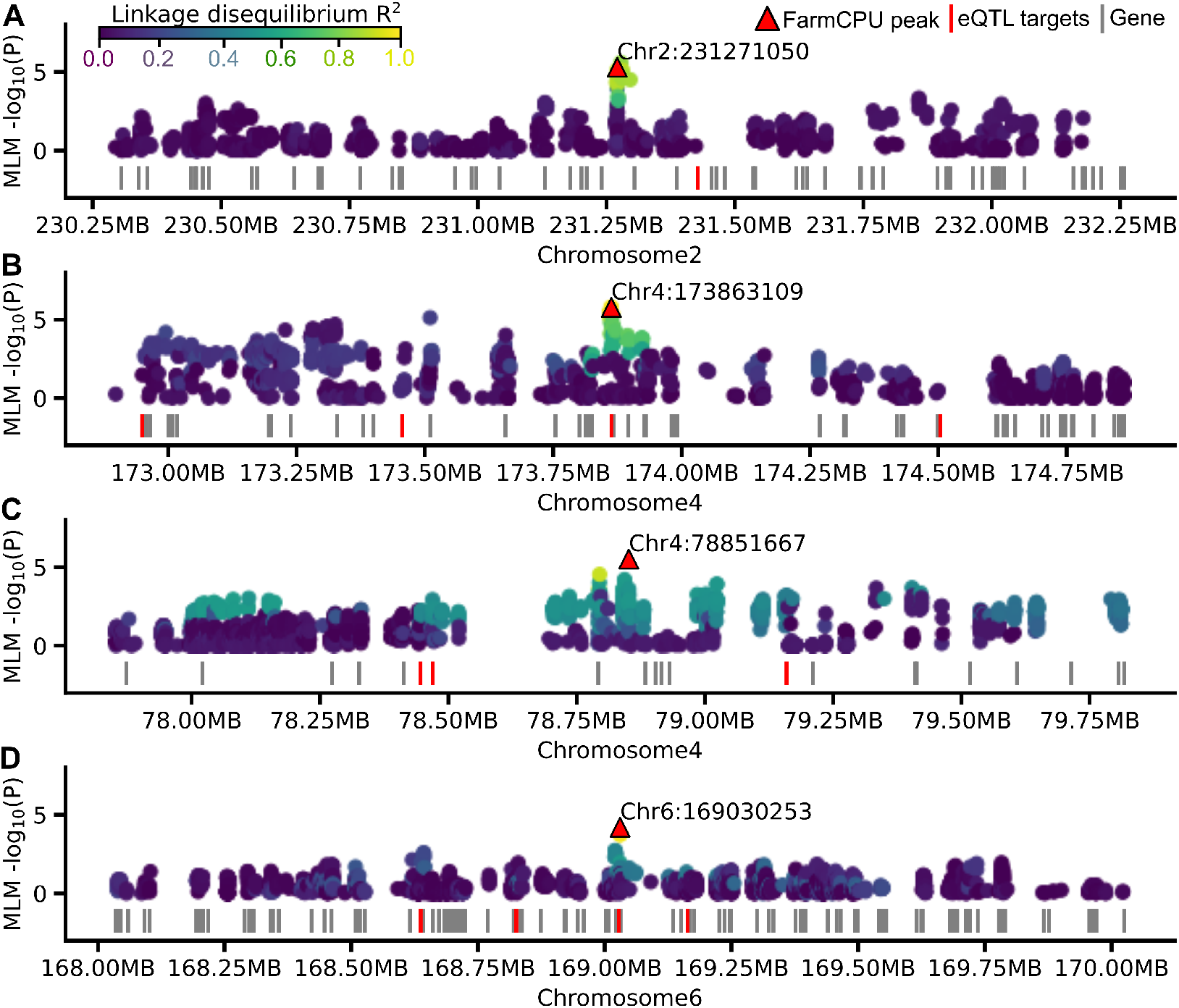
Candidate genes associated with each of four significantly trait associated SNPs. Local manhattan plots showing the statistical significance of markers within one megabase up or downstream of each of four significant trait associated SNPs (A-D). Y-axes indicate the statistical significance assigned to each marker (circles) by the GEMMA univariate linear mixed model^25^. Color coding of circles indicates the degree of linkage disequilibrium observed between a given marker and the trait associated marker. Vertical bars indicate the positions of annotated gene models. Vertical bars in red indicate annotated gene models where GWAS for the expression level of that gene in the same population identified either the trait associated SNP or a SNP in ≥ 0.8 LD with the trait associated SNP.

### Variation in southern corn rust disease severity is associated with variation in seedling transcript abundance

Transcriptome-wide association studies can play a complementary role to genome wide studies by identifying genes which play important roles in determining phenotypic outcomes which are missed by the latter analysis^33,34^. Expression level data were generated for different subsets of the Wisconsin Diversity Panel as part of two published studies^21,24^. While batch effects exist between the two datasets (Figure S5), both sets were incorporated into the transcriptome wide association study with five expression level principal components incorporated as controls alongside five principal components calculated from genetic marker data (see Methods). The expression level of a single gene, Zm00001d035185, a RNAse E/G-like protein (RNE) located on chromosome 6 at 8.6 MB, was significantly associated with variation in southern rust disease severity after correcting for multiple testing employing the Benjamini–Hochberg procedure (fdr < 0.05) (Figure S6A). Published expression datasets indicated that this gene was highly expressed in the leaves of maize throughout its entire lifecycle (Figure S7A). The Arabidopsis homolog of Zm00001d035185, AT4G37920, exhibits high expression in leaves and the protein product was predicted to be localized to guard cells (Figure S7B). Guard cells are known to be involved in pathogen recognition and immune response suggesting a plausible mechanism for the link between expression of this gene and southern corn rust disease severity^35,36^. The next four most significantly associated genes including Zm00001d050928 (p = 0.000021) Zm00001d035195 (p = 0.000021), Zm00001d039975 (p = 0.000025) and Zm00001d013049 (p = 0.000026) were also reported to be related to plant immune response to pathogen infections (see discussion) (Figure S6B-E).

## Discussion

In this study we took advantage of an unanticipated southern corn rust outbreak in eastern Nebraska during the 2020 growing season to assess the susceptibility or resistance of a set of 689 genotypes drawn from the Wisconsin Diversity Panel which were grown as part of a preexisting experiment and had neither lodged nor senescensed at the time of the outbreak. As previously reported and studied, different individuals scoring disease severity using a common ordinal scoring metric generated differing distributions of disease severity scores and different values even when scoring the same plots^37^. Controlling for variability in the medians and distributions of assigned values between scorers substantially increased the overall heritability of the southern rust disease severity trait and enabled the identification of four statistically significant trait associated SNPs and one statistically significant trait associated transcript. In recognition of the history of maize QTL being associated with upstream regulatory regions in only partial linkage with the coding sequence of the target gene, expression GWAS was used to identify sets of candidate genes associated with each individual trait associated SNP.

The only gene Zm00001d007424 linked to the trait associated SNP on chromosome 2 via expression level variation is annotated to encode a Long chain base biosynthesis protein 1 (LCB1) that along with LCB2 forms the heterodimer serine palmitoyltransferase (SPT) that catalizes the first rate-limiting step of *de novo* sphingolipid synthesis (Figure 4A). Loss of LCB1 lead to abolishment of programmed cell death initiated by the production of reactive oxygen intermediates (ROIs)^38^. There is a growing body of evidence supporting this sphingolipid mediated programmed cell death is associated with plant immunity to pathogen infection such as the maize pathogen *Fusarium verticillioides* and the tomato pathogen *Alternaria alternata f. sp. lycopersici* as reviewed by Berkey *et al*., 2012^39^).

Among the three genes (Zm00001d050283, Zm00001d050284, Zm00001d050293) linked to the first trait associated SNP on chromosome 4 (79 MB) via expression level variation, Zm00001d050283 is annotated as Sulfate transporter 1.2, this sulfate transporter is highly conserved across kingdoms and its homology in tomato is highly expressed upon vascular pathogen *Verticillium dahliae* infection^40^. Zm00001d050284 is annotated as tRNA modification GTPase (trmE) of which the function plays pivotal roles in protein translation in plants. A natural loss of function variant allele of the rice PLEIOTROPIC DEVELOPMENTAL DEFECTS (PDD) which encodes the same E family tRNA modification GTPase lead to reduction of ribosome biogenesis and proteins related to photosynthesis which could affect the leaf fitness^41^; and Zm00001d050293 encodes the trehalose-6-phosphate synthase 9 (TRPS9), a key enzyme involved in trehalose-6-phosphate metabolism, is involved in plant resilience^42^ and immunity response to pathogen attacks in tomato^43^.

Among the four genes (Zm00001d051869 (no annotation), Zm00001d051884, Zm00001d051893, Zm00001d051914 (no annotation)) linked to the second trait associated SNP on chromosome 4 (174 MB) identified via expression GWAS, Zm00001d051884 encodes a hydroxyproline-rich glycoprotein family protein; Zm00001d051893 encodes a TPR-like superfamily proteins which have been reported to be involved in disease response in plants. For example, a loss of function allele of rice TRP-domain RNA-binding protein BSR-K1, bsr-k1 rendered a broad spectrum disease resistance in rice^44^. The potential mechanisms of this protein family conferring disease resistance is likely hormone responsive. Noteably, this is the only gene which was also linked to the chromosome 4 (174 MB) trait associated SNP directly, the SNP is located within the gene, or through the presence of SNPs in high LD with the trait associated SNP within the gene body (Figure 4 B).

Among the four genes (Zm00001d039004, Zm00001d0390020(no annotation), Zm00001d039039, Zm00001d039043) linked to the trait associated SNP on chromosome 6 via either expression level variation or LD, NAC87(Zm00001d039004) is a NAC-Type Transcription Factor that has been identified to positively promote modulate Reactive oxygen species accumulation, hyper sensitive response and cell death in response to stresses in oilseed rape (*Brassica napus L*.)^45^. As a transcription factor, its downstream components included RbohB which is involved in ROS production, VPE1a and ZEN1 which regulate cell death, WRKY6 and ZAT12 which are involved in leaf senescence and commonly known plant disease resistant genes like PR2 and PR5^45^. Zm00001d039039 encodes a chloroplastic Cold-regulated 413 inner membrane protein 2; Aspartic proteinase A1 (Zm00001d039043) is also likely involved in disease resistance. In arabidopsis, an extracellular aspartic protese encoded by CDR1 lead to dwarfing and resistance to virulent *Pseudomonas syringae* by activating micro-oxidative bursts and defense-related gene expression in a salicylic-acid-dependent manner^46^.

In addition, the other four of the top five most significantly associated genes, while not individually passing stringent multiple testing correction, includes several intriguing potential links to disease resistance. In order of decreasing statistical significance they are: Zm00001d050928 (p = 0.000021), a gene with no functional annotations (Figure S6B), Zm00001d035195 (p = 0.000021), a gene encoding a kysine-specific histone demethylase 1 (LSD1) that is reported to be involved in plant growth, flowering time, hormone response, stress response and circadian regulation^47–49^ (Figure S6C), Zm00001d039975 (p = 0.000025), a gene with no functional annotations (Figure S6D), and Zm00001d013049 (p = 0.000026), a gene encoding a exocyst complex component (EXO84B), for which knockout alleles of the arabidopsis homolog have been shown to exhibit significantly increased susceptible to the hemi-biotrophic pathogen *Phytophthora infestans* infection^50^ (Figure S6E). However, given the lack of statistical significance after correcting for multiple testing correction, it must be kept in mind that one, or more, or all, of these genes may represent false positive associations. Southern corn rust have long been primarily considered a threat to maize production in tropical and subtropical climates. However, forecast changes in climate are likely to increase the frequency and severity of southern corn rust outbreaks in more temperate production regions^2^. While a major resistance locus, *RPP9* was incorporated into a number of commercial maize hybrids in the USA^11^, breakdowns of the resistance to southern corn rust conveyed by this locus began to be reported over one decade ago^1,13^. While qualitative resistance loci have been largely sourced from tropical maize germplasm^1^, here we show that significant quantitative resistance exists and can be mapped to specific loci within a set of 689 diverse temperate adapted maize inbred lines. Given the potential for yield losses of 39-50% from severe and early season southern rust outbreaks and the increasing potential for southern rust outbreaks, developing phenotyping, genetics, and breeding efforts to ameliorate or at least mitigate potential losses from such an outbreak are clearly needed.

## Materials and methods

### Field experiment and phenotyping

A set of 752 maize genotypes drawn from the set of 942 genotypes included in the Wisconsin Diversity Panel^21^ as well as integrated checks were planted at the Havelock Farm research facility of the University of Nebraska-Lincoln on May 6^th^, 2020 (40.852 N, 96.616 W). Each plot consisted of two rows of the same genotype, and was 7.5 foot long with 30 inch row spacing and 2.5 foot alleyways between sequential plots. Significant southern rust infection was observed on August 14^th^, 2020. Scoring for southern corn rust disease severity was conducted on August 19^th^, 2020. Severity of rust was assessed by three individuals using a five point rating scale with 0.0 being completely free of southern rust and 4.0 being the most susceptible^27^. In order to assess and correct for individual specific biases in scoring, a subset of 284 plots was selected and scored by all three individuals (Supplementary Document 1). The plots that died, desiccated or lodged were skipped for disease severity scoring.

Using the row and column coordinate of the control plots and southern rust disease severity by P1 and P3 as independent variable, disease severity scored by P3 as dependent variable, we performed Ordinary Least Square Regression (OLS) implemented in statsmodels in python (v 3.8) to calculate the intercepts and coefficients of row, column and disease scores of the regression line that used to predict P3 disease score by the corresponding dependent variables of all other plots (Supplemental table 1 & 2). The resulted scoring was then used for calculating genotype BLUPs for disease severity using the lme4 (v 1.1-26) package in R (v 4.0.2)^51^, genotype and scorers were fitted in the model as one of the random effects, model = MeanScore (1 |Genotype) + (1 |Scorer). After correction by regression, the ‘Scorer’ variable explained less than one half of one percent of total variance.

### Genotype dataset

High density genotyping was conducted using a set of 23,672,341 *a priori* segregating genetic markers from the Maize HapMap3 dataset^22^. A set of 129 lines overlapped between the HapMap3 dataset and the 942 lines of the Wisconsin diversity panel. For the remaining lines, *a priori* were scored using published resequencing data (452 genotypes were taken from Connor *et al*., 2020^23^) or published RNA-seq data (361 lines taken from Mazaheri *et al*., 2019^21^ or Hirsch *et al*., 2014^24^).

Sequence data were first trimmed using using Trimmomatic(v 0.33) with parameter settings “-phred33 LEADING:3 TRAILING:3 slidingwindow:4:15 MINLEN:36 ILLUMINACLIP:TruSeq3-PE.fa:2:30:10“^52^. The resulting trimmed readers were mapped to the B73 RefGen_v4 maize genome assembly^53,54^ using BWA-MEM (v 0.7) with default parameter settings^55^. Duplicated reads resulted from PCR amplification were marked using picard (v 2.22)^56^. The resulting BAM files were used to call genotypes for the 23,672,341 *a priori* SNP sites using GATK toolkit(v 5.1)^57^. After genotyping, individuals with missing data were imputed with Beagle/5.01 with parameter settings ‘window=1 overlap=0.1 ne=1200’ with the entire HapMap3 population employed as a reference panel^58^.

Hapmap3 includes 83M total variants^22^. The subset of 23M *a priori* SNP sites employed above was created by filtering the unimputed hapmap3 dataset downloadable from MaizeGDB^59^. First individuals with missing data rates higher than 0.6 or the inbreeding coefficient slower than 0.9 were removed. Second, individual markers with minor allele frequencies <0.01 or missing data rates >0.6 among the remaining individuals were removed. Then the remaining markers were inmputed using Beagle/5.1 with parameter settings ‘window=1 overlap=0.1 ne=1200’^58^. Finally, sites with heterozygosity rates >0.2 after imputation were removed, resulting in the final *a priori* segregating SNP dataset.

### Genome-wide Association Analyses

For genome wide association studies, the initial SNP set was further filtered through the removal of SNPs with a minor allele frequency *leq* 0.1 and SNPs with a heterozygous genotype call frequency *geq* 0.01 across the 689 individuals, resulting in a set of 2,730,913 SNPs. Kinship matrices and the first five population structure principal components were calculated from unlinked makers (R2 ≤ 0.2) using PLINK (v 1.9)^60^. Two separate GWAS algorithms were employed. First, GWAS was conducted using the univariate mixed linear model (LMM) implemented in GEMMA (v 0.98.3) using the Kinship matrices and the first five principal components calculated from marker data to control the confounding effects of the population structure^25^. Secondly, GWAS was conducted using the FarmCPU algorithm^61^ as implemented in rMVP (v 1.0.4)^62^. In order to estimate the stability of significant trait associated SNPs identified by FarmCPU, a resampling strategy was employed with the analysis run 100 times each with a different randomly selected subset of the 689 genotypes for which phenotypic and genotypic data was available^63,64^. Three population structure principal components were calculated from the genotype data of the sampled population to control the population structure. Individual SNPs which were statistically significantly linked to the trait in question in ≥ 10 of 100 resampling GWAS were considered both statistically significant and stable trait associated SNPs.

### Expression quantitative trait loci mapping

Published expression values (FPKM) for annotated maize genes across the 689 genotypes with southern rust disease severity scores were retrieved from previous studies^21,24,65^. These values were normalized via a Box-Cox transformation as described in^66^. matrixEQTL was used to conduct GWAS using the same set of SNP markers described above, and the normalized expression levels of each gene located within 1 megabase up or downstream of a trait associated SNP as individual molecular phenotypes^67^. The matrixEQTL analysis incorporated 5 PCs used for GWAS previously in this study as covariates.

### Transcriptome wide association study

Tests for associations between transcript levels and disease severity scores were conducted for each of a set of 21,820 genes which were expressed at levels >1 FPKM in at least 345 of the 689 individuals scored for southern rust disease severity. Gene expression levels were treated as explanatory variables for southern corn rust severity score BLUPs. The association study was conducted by fitting the gene expression level as a random effect, 5 genetic PCs and 5 expression PCs as fixed effects to control the population structures using lme4 (v 1.1-26) package in R (v 4.0.2)^51^. P values of genotype coefficients for each gene expression-disease severity score BLUP pair were ranked as references of association significance.

## Supporting information

Raw southern rust disease severity scores

## 1 Data availability statement

Two sets of RNA-seq data used in this study are from Mazaheri *et al*., 2019^21^ available at NCBI BioProject accession PRJNA189400 and Hirsch *et al*., 2014^24^ available at NCBI BioProject accession PRJNA437324 respectively. Resequencing genome sequencing data are from O’Connor *et al*., 2020^23^ available at NCBI BioProject accession PRJNA661271. The maize hapmap3 genomic data are from Bukowski *et al*. (2018)^22^ and available from https://datacommons.cyverse.org/browse/iplant/home/shared/commons_repo/curated/Qi_Sun_Zea_mays_haplotype_map_2018/282_onHmp321. The SNP dataset generated for genome wide association in this study is openly available in figshare at https://figshare.com/articles/dataset/WiDiv689_maf1het01_vcf_gz/14365544.

## 2 Acknowledgements

This work was supported by the National Science Foundation under grant OIA-1557417 to JCS, the Foundation for Food and Agriculture Research under award number – Grant ID: 602757 to JCS, and the US Department of Energy Advanced Research Projects Agency-Energy (ARPA-E) under Contract No. 18/CJ000/01/0 This project was completed utilizing the Holland Computing Center of the University of Nebraska, which receives support from the Nebraska Research Initiative. We thank Dr. Marcin Grzybowski for the technical supports of this work.

## 3 Author contribution

J. C. S., G. S., and R. V. M., conceived this research. J. C. S., G. S., and R. V. M., designed and directed the study. G. S., R. V.M., and J. D. T., scored the disease severity. G. S., and R. V. M., performed data processing and analysis. G. S., R. V. M., and J.C. S., drafted the manuscript. The final version of the manuscript was generated with input and contributions from J. D. T. All authors approved the final version of the manuscript.

---

**Table S1.** Southern rust scored by this study and Medina *et al*., 2007^28^

| Plot | Genotype | Mean score | Blup | Medina *et al* 2007 |
| --- | --- | --- | --- | --- |
| 2177 | Va59 | 0 | 0.857557494 | 0 |
| 2219 | INBRED 2-687 | 0 | 1.125114209 | 0 |
| 2633 | INB 101LFY/LFY (A632 X M16 S5) | 0 | 1.61051951 | 1 |
| 2712 | Ky228 | 0 | 1.047181909 | 1 |
| 1641 | C103 | 1 | 0.9082285 | 1 |
| 2765 | Mo17 | 1.5 | 1.707236075 | 1 |

**Table S2.**
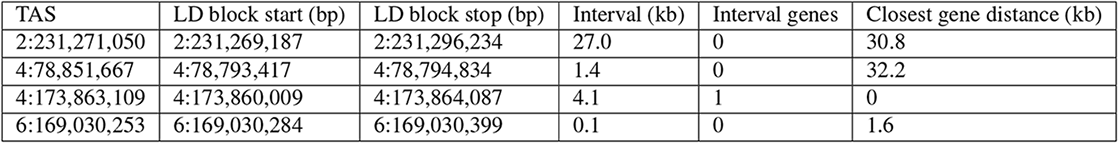
Local linkage disequilibrium for four FarmCPU peaks

**Figure S1.**
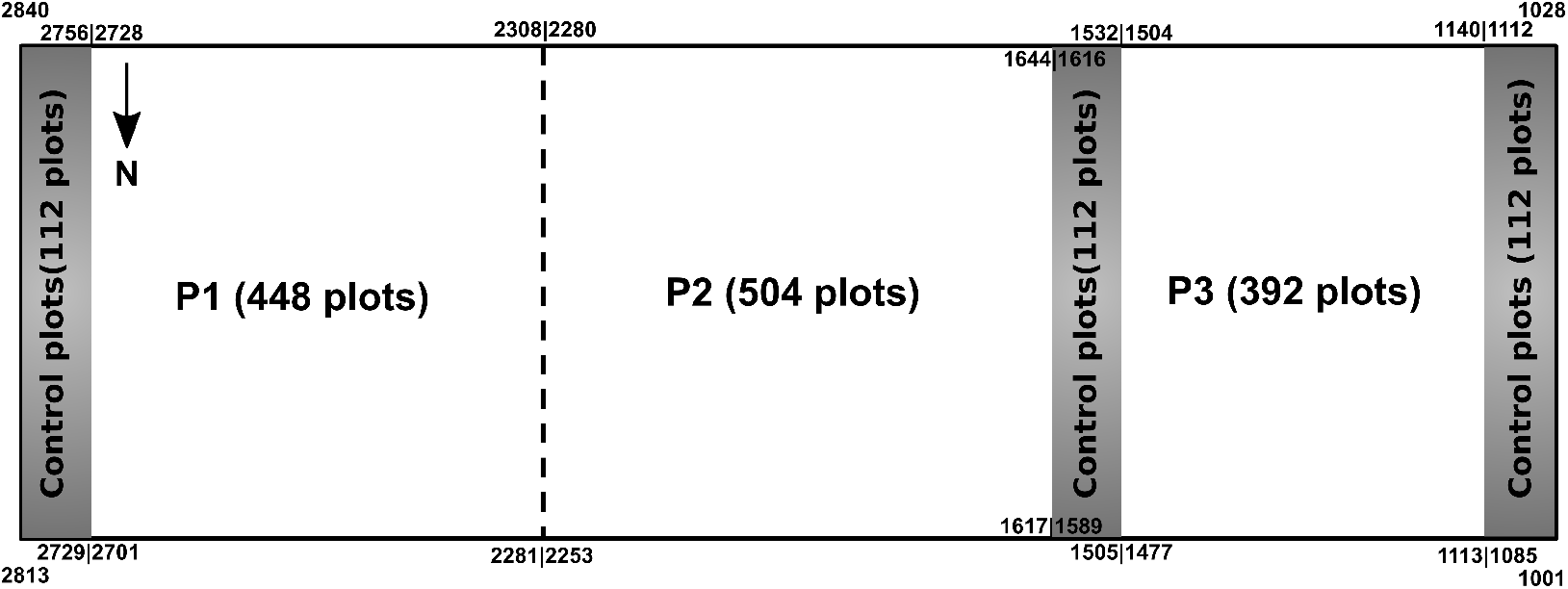
Field layout and the regions assigned to scorer P1, P2 and P3.

**Figure S2.**
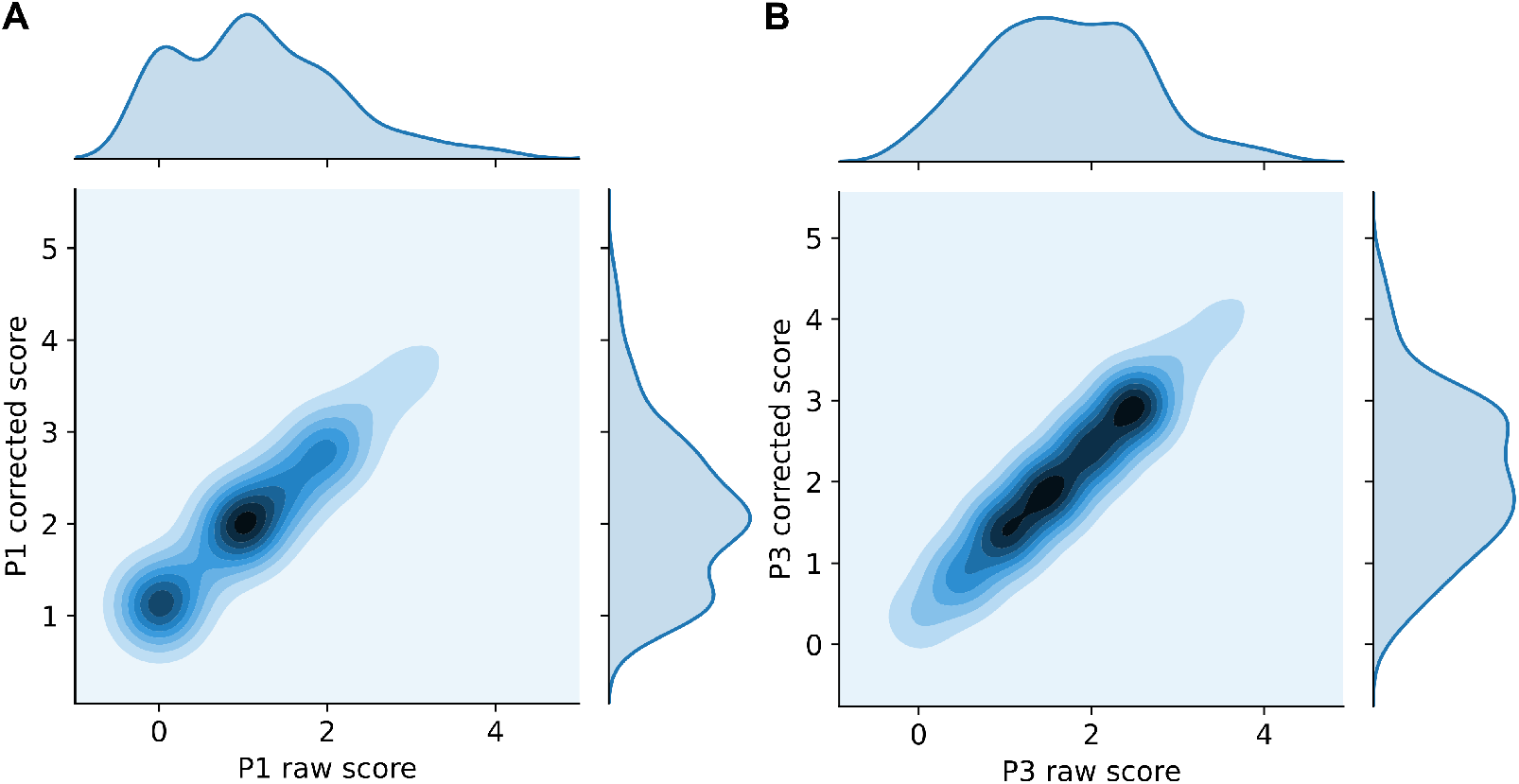
Score correction via linear regression. A) Ordinary Least Square Regression and the predicted score of disease severity scored by P1 using P2 as dependent variable. B) Ordinary Least Square Regression and the predicted score of disease severity scored by P3 using P2 as dependent variable.

**Figure S3.**
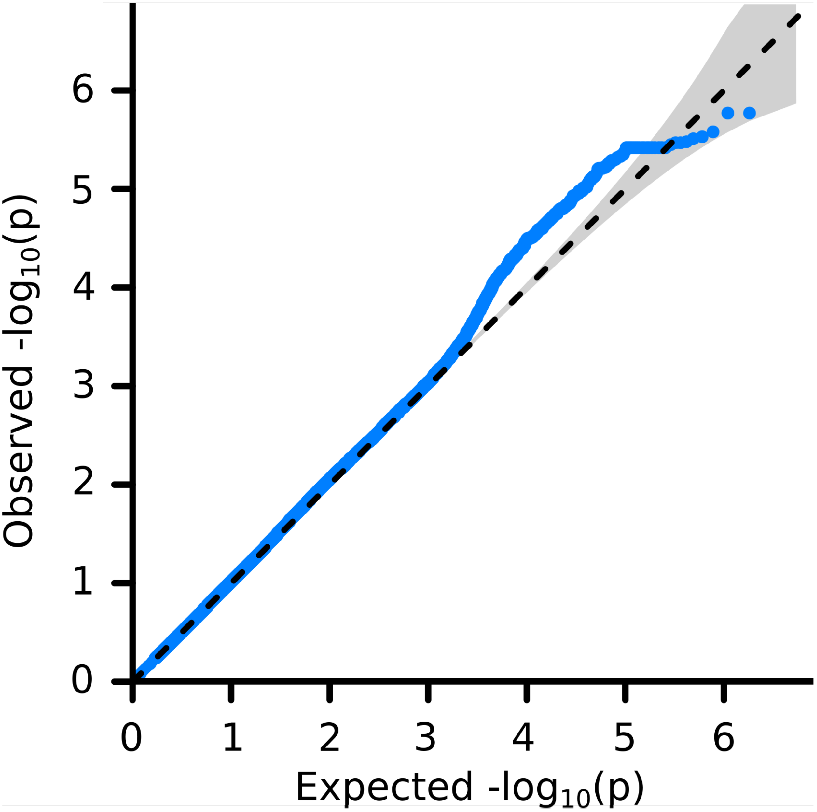
Quantile-Quantile plot of uncorrected p values calculated by the Mixed Linear Model implemented in GEMMA. The −log 10 transformed observed P values were plotted against their expected values under the null hypothesis that the markers have no effect (the main diagonal line).

**Figure S4.**
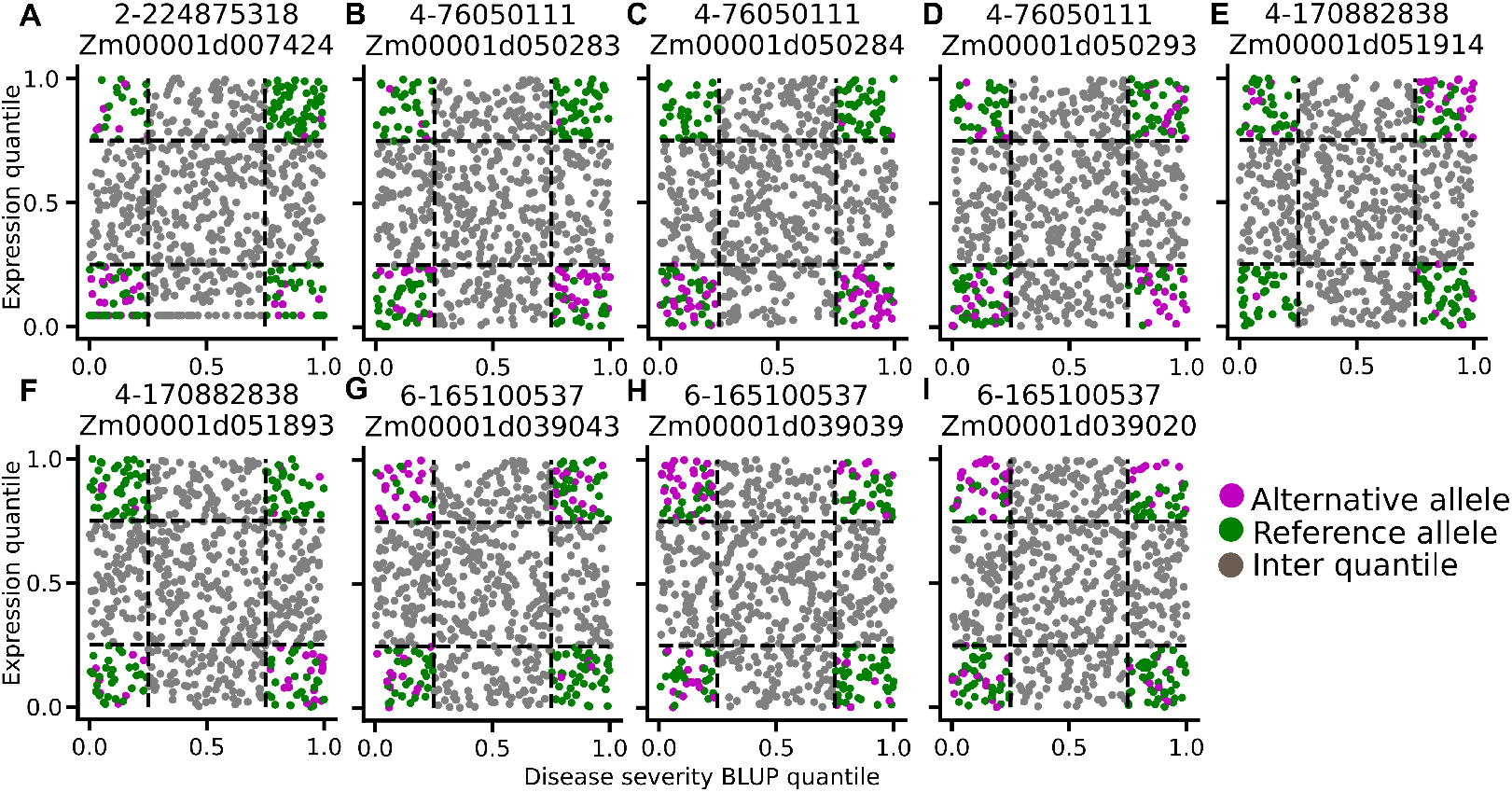
Correlation between the expression of eQTL targets and disease severity scores discriminated by the alleles of eQTLs. Percentiles of disease severity score BLUPs (x-axis) and the percentiles of eQTL targets expression (y-axis) for all of the individuals carrying homozygous eQTLs were plotted, the inter quantile (0.25-0.75) for expression level and disease severity score BLUPs are plotted in grey. Only in the 0-0.25 and 0.75-1.0 quantiles are the points color coded by alleles.

**Figure S5.**
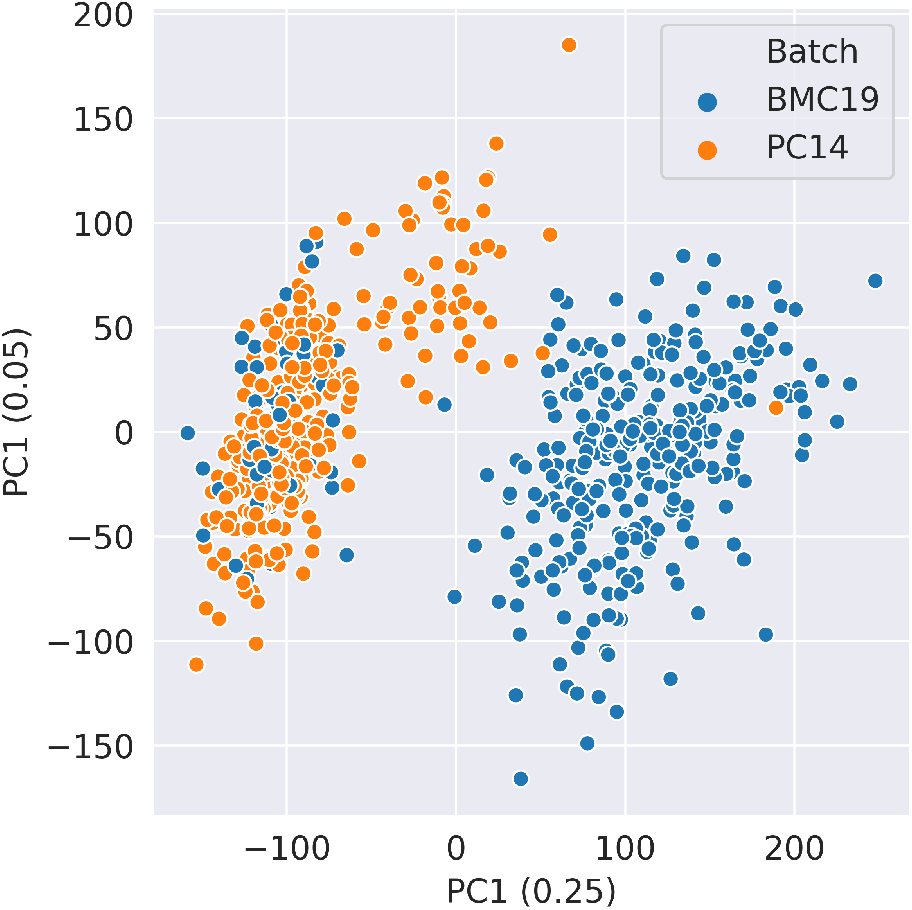
Principal components calculated from the transcriptomics data for 689 individuals. The expression data generated from two different batches “BMC19” by Mazaheri *et al*., 2019^21^ and “PC14” by Hirsch *et al*., 2014^24^ were color coded. PC1 which explained 25% of the expression variation well captured batch effects.

**Figure S6.**
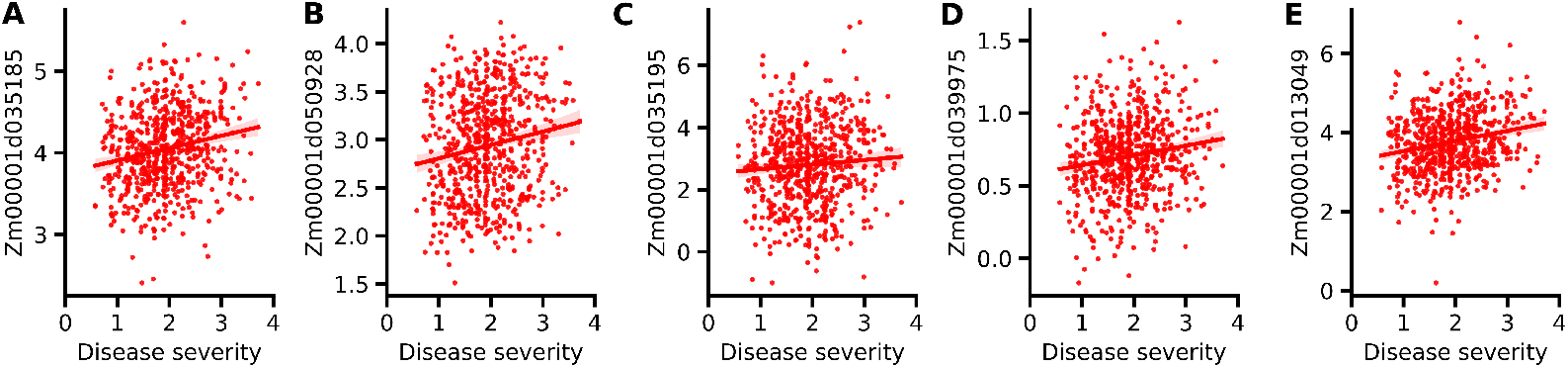
Candidate genes involved in southern corn rust susceptibility or resistance identified via transcriptome wide association. The relationships between disease severity score BLUPs and seedling expression levels for five most significantly associated genes identified by TWAS (A-E)

**Figure S7.**
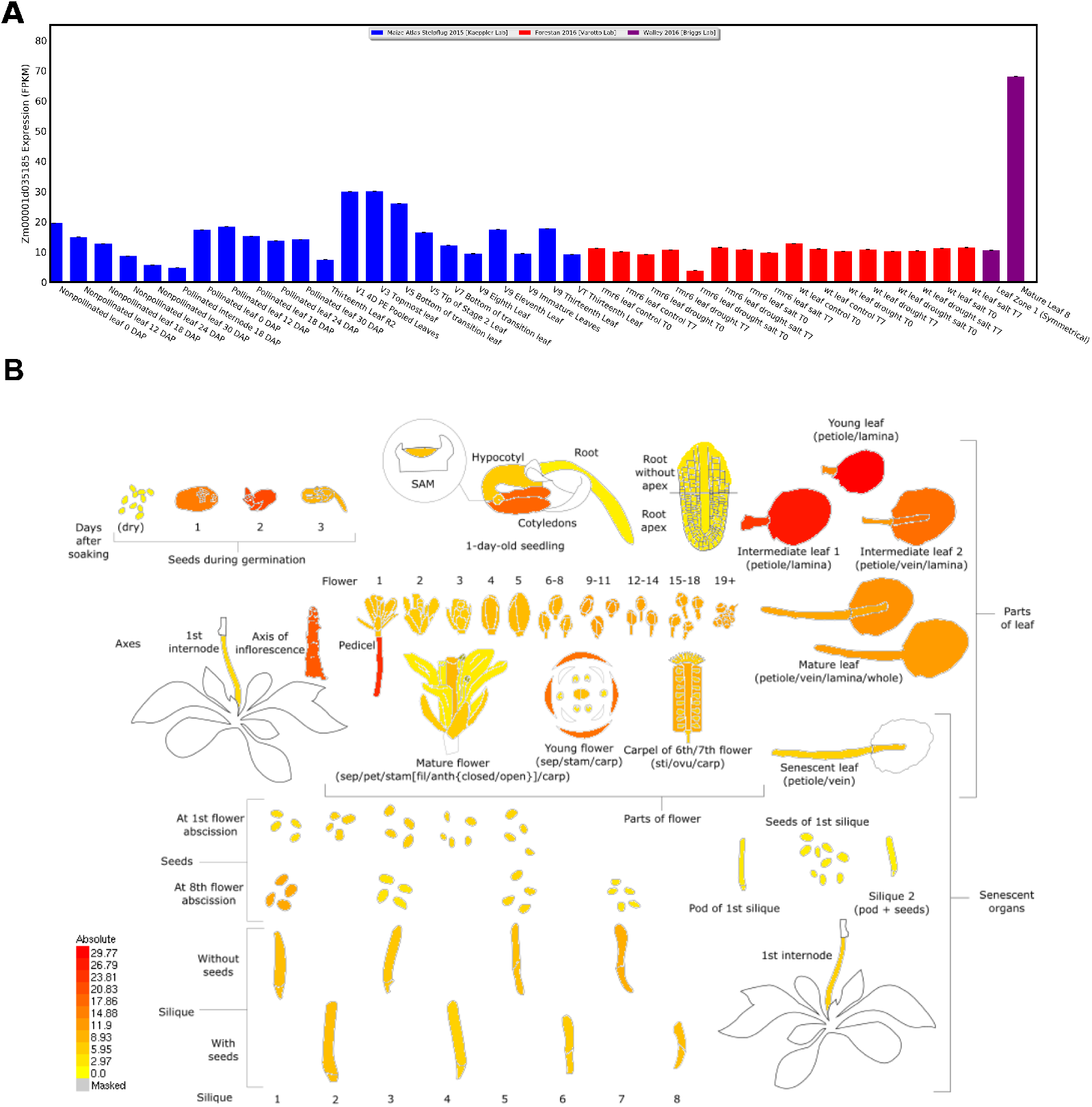
The expression pattern of RNAse E/G-like protein (RNE) in maize (A) and Arabidopsis (B)

